## Supplementary Materials for "Temporal control of gene expression by the pioneer factor Zelda through transient interactions in hubs"

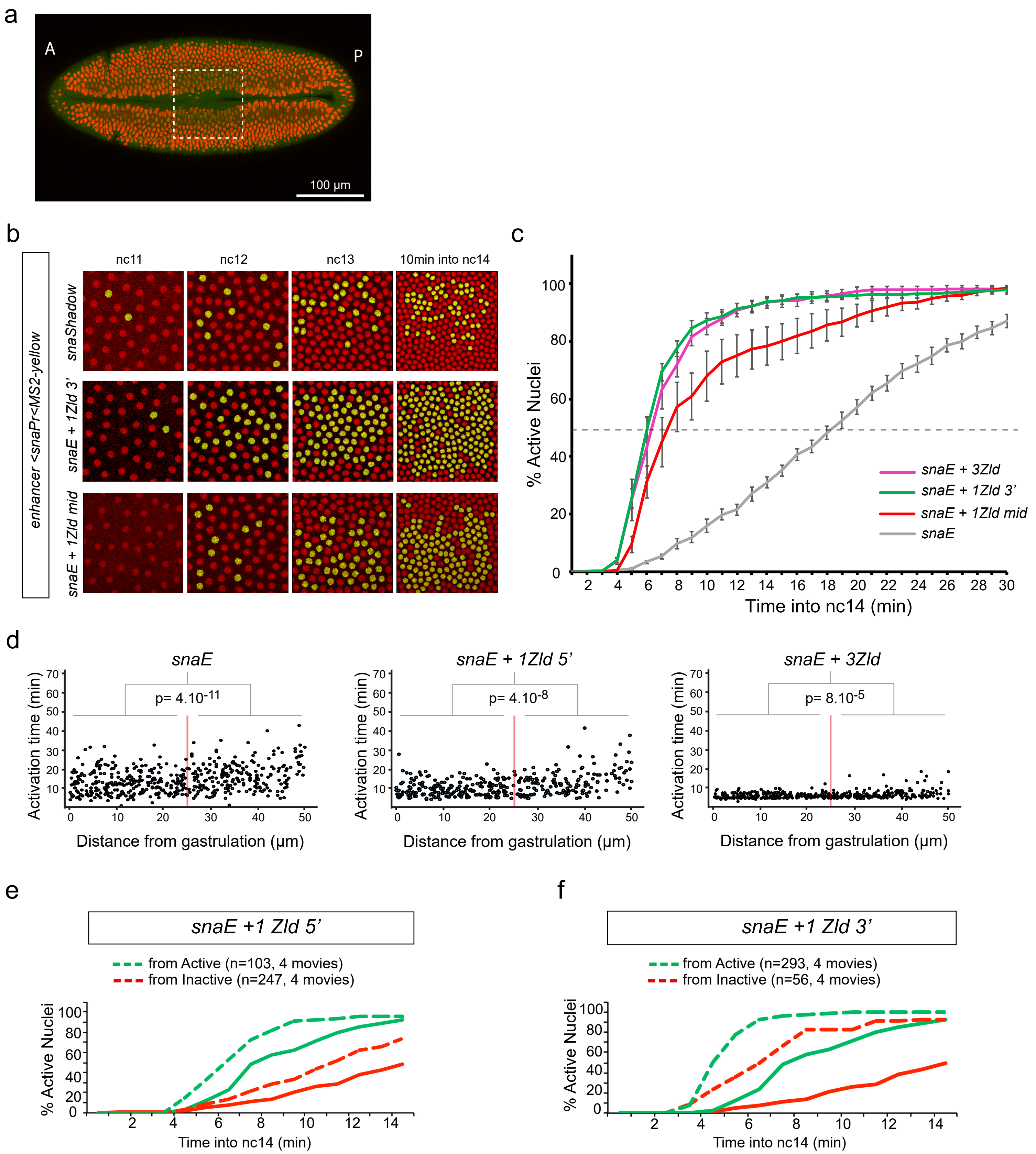


**Supplementary Figure 1: (related to Figure 1)**

(**a**) Maximum intensity projected Z-stacks of a representative gastrulating embryo tile scan showing the imaged region (dashed line) related to Figure 1. Nuclei are in red and MS2 dots in green. The embryo is oriented with Anterior (A) to the left, and Posterior (P) to the right.

(**b**) Snapshots of maximum intensity projected Z-stacks from live-imaging movies of transcriptional activity driven by *sna-shadow*, *snaE+1Zld 3'* and *snaE+1Zld mid* transgenes from nc11 to nc14. Nuclei exhibiting transcriptional activity were false-colored in yellow. (**c**) Synchrony curve for *snaE+1Zld 3'* (green line), *snaE+1Zld mid* (red line), *snaE+3Zld* (purple line) transgenes compared to that of the *snaE* (grey line) transgene. Statistics: *snaE* (6 movies, n=970 nuclei), *sna-shadow* (4 movies, n=612 nuclei), *snaE+3Zld* (4 movies, n=824 nuclei), *snaE+1zld* mid (4 movies, n=659 nuclei). Error bars represent SEM.

(**d**) Scatter plot of the activation time as a function of the distance from the ventral furrow at nc14, for *snaE* (left panel), *for snaE+1Zld5'* (middle panel) and for *snaE+3Zld* (right panel) transgenic embryos. Statistics: *snaE* (6 movies, n=490 nuclei), *snaE+1Zld5'* (4 movies, n=350 nuclei), *snaE+3Zld* (4 movies, n=411 nuclei). Comparison of the timing of activation in nc14 for nuclei located 0-25µm around the ventral furrow and those located more distally at 25-50µm. P-values using a two-tailed student t-test are shown.

(**e-f**) Quantification of transcriptional memory in *snaE* (**e-f**) *snaE+1Zld 5'* (**e**) and *snaE+1Zld 3'* (F) transgenic embryos. Temporal dynamics from *snaE* are shown as solid curves, while those from extra Zld binding sites transgenes are represented with dashed curves (**e-f**). Each graph shows the percentage of transcriptionally active nuclei in the first 15min of nc14 for nuclei coming from active nc13 mothers (green curves) and inactive nc13 mothers (red curves). Only nuclei located 50µm around the gastrulation line are considered.

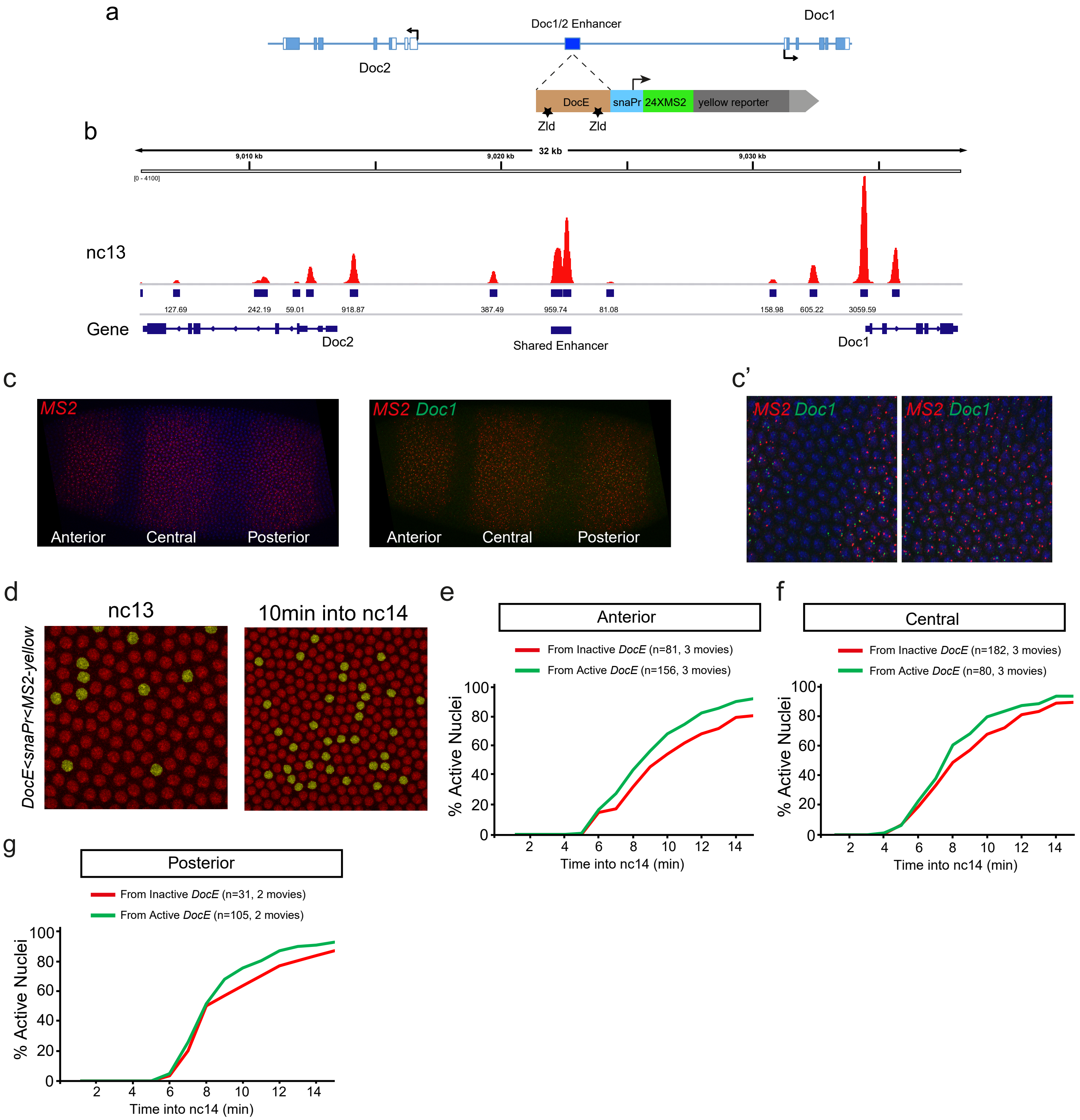


**Supplementary Figure 2: (related to Figure 2)**

(**a**) Schematic representation of the *Doc1/2* locus and localization of the regulatory element (*Doc-Enhancer*) used in this Figure. (**b**) The *Doc-Enhancer* was designed according to Zelda ChIP data on nc13 embryos[1]. A transgenic line was created with the *Doc-Enhancer* driving expression of an *MS2-yellow* reporter from a *sna* minimal promoter (*DocE transgene*) (**a**).
(**c**) A nc14 *DocE* transgenic embryo was hybridized with a probe against *MS2* (red) and an intronic probe against endogenous *Doc1* (green). Maximum intensity projected Z-stacks confocal images with MS2 signal alone (left panel) or merged with *Doc1* (right panel). The *DocE* transgene drives expression in the dorsal mesoderm in a tripartite pattern (anterior, central and posterior) (**c'**) Zoomed images from the embryo shown in (**c**). (**d**) Snapshots from a maximum intensity projected Z-stack movie of a *DocE* transgenic embryo, imaged in the central dorsal ectoderm domain. Nuclei exhibiting an active transcriptional dot are false colored in yellow. The *DocE* transgene leads to a stochastic expression in nc13, which allows transcriptionally active mothers to be distinguished from their inactive neighbors.
(**e-g**) Quantification of transcriptional memory in *DocE* transgenic embryos in three regions of the dorsal ectoderm, anterior (3 movies, n=237 nuclei) (**e**), central (3 movies, n=262 nuclei) (**f**) and posterior (2 movies, n=136 nuclei) (**g**).

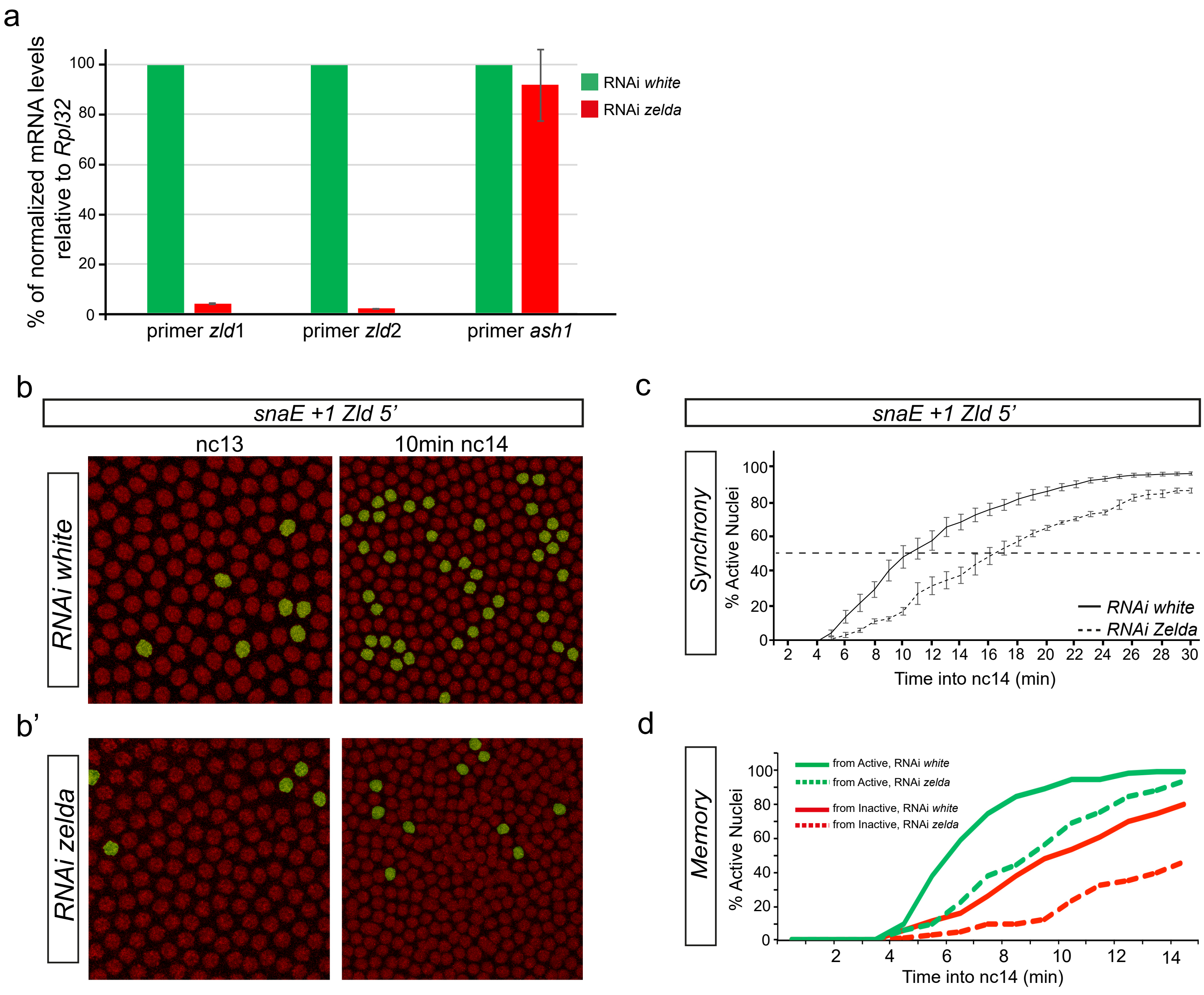


**Supplementary Figure 3: (related to Figure 3)**

(**a**) mRNA quantification using qRT-PCR in *white-RNAi* and *zld-RNAi* 0-2-hr embryos. *RpL32* was used as a control mRNA for normalization, *ash1* was used as a control mRNA. The levels of mRNA for *zld* and *ash1* in *zld-RNAi* embryonic extracts (red) were normalized to their levels in *white-RNAi* extracts to 100% (green). For each genotype, experiments were performed on four biological replicates. Two primer pairs were used to quantify *zld* mRNA (zld1 and zld2). Error bars represent SEM. (**b, b'**) Snapshots of maximum intensity projected Z-stacks *Drosophila* embryo movies expressing the *snaE+1Zld5'* transgene at nc13 or at nc14 in a *white-RNAi* (**b**) or in a *zld-RNAi* genetic background (**b'**). Nuclei with active transcription spots are false colored in yellow.
(**c**) Temporal coordination for the *snaE+1Zld5'* transgene in a *white-RNAi* (solid curves) and in a *zld-RNAi* genetic background (dashed curves). Statistics: *SnaE+1Zld5' white-RNAi* (5 movies, n=452 nuclei) and *snaE+1Zld5' zld-RNAi* (5 movies n=167 nuclei). (**d**) Kinetics of activation during the first 15min of nc14 driven by the *snaE+1Zld5'* transgene in *white-RNAi* (solid curves) or *zld-RNAi* embryos (dashed curves). The kinetic of nuclei coming from transcriptionally active mother nuclei in nc13 (green curves) is compared to those arising from inactive mothers (red curves). In Supplementary Fig. 3, only nuclei located within a 25µm rectangle from 'pseudo ventral furrow' are considered.

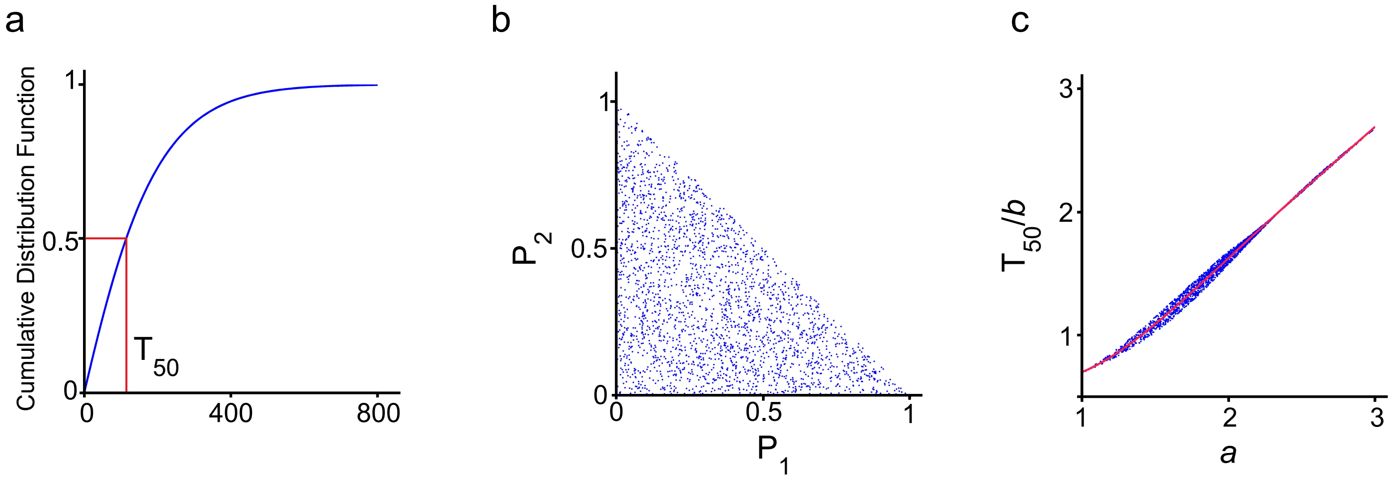


**Supplementary Figure 4: (related to Figure 4)**

(**a-c**) Graphs resulting from simulated data from the homogeneous jump model. (**a**) We defined T50 as the time needed to reach 50% of total activation; the inverse of this time is thus a measure of the speed of reactivation. Using the homogeneous jump model, we computed the random component of this time for many values of *'b'* and for many values of p1 and p2, (probabilities to reach activation in one and two jumps, respectively).

(**b**) Panel showing the uniformly distributed values of p1 and p2. (**c**) When projected onto the plane of the variables *'a'* and T50/*b* (blue dots), the simulated data appear close to the curve of equation:T50/*b*= 0.085a4−0.7834a3+2.651a2−2.814a+1.56 (red line).

These findings imply that the activation time (T50) is proportional to *'b'* and increases with *'a'*. Furthermore, in units of *'b'*, this time depends on *'a'* only.

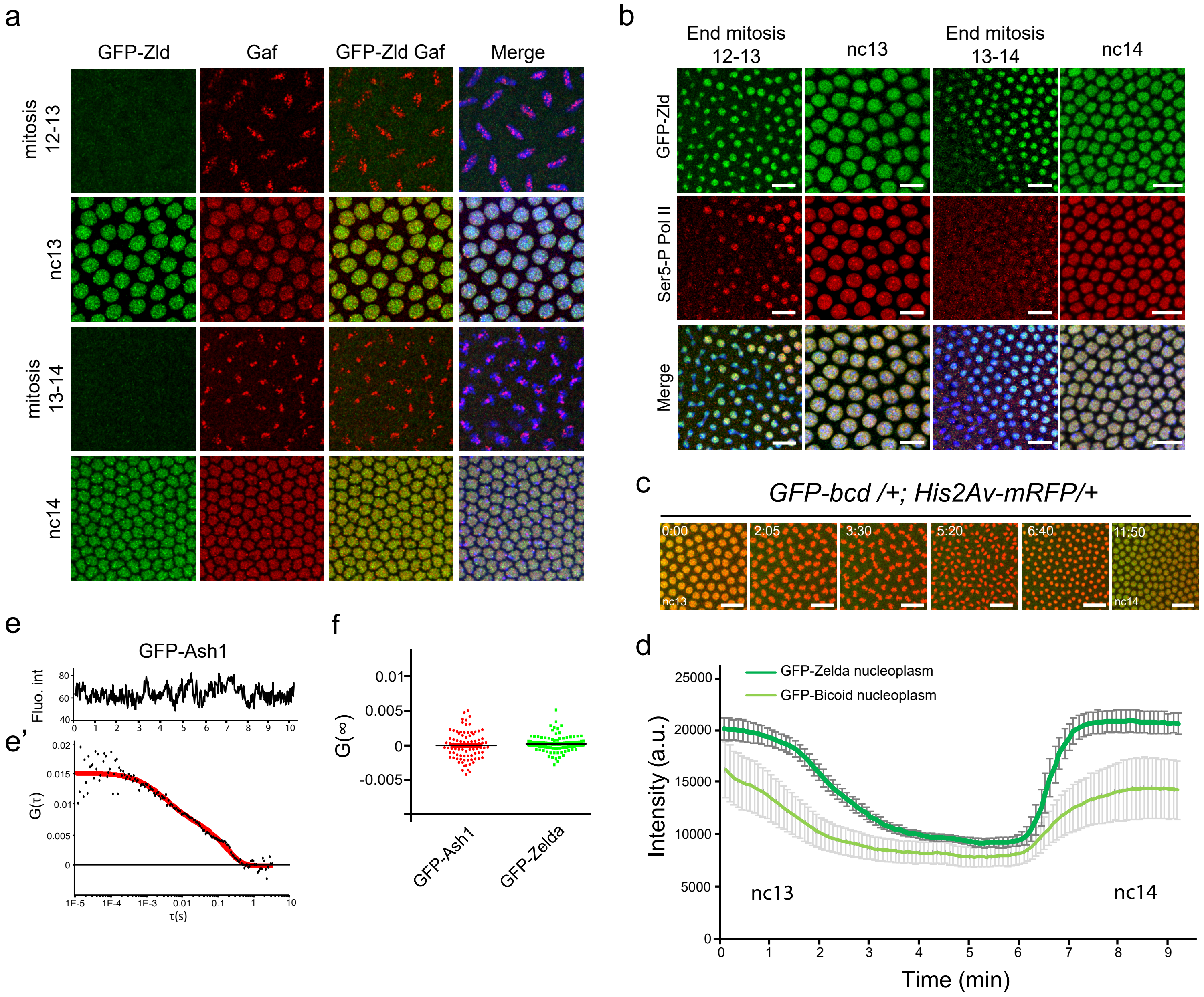


**Supplementary Figure 5: (related to Figure 5)**

(**a**) Contrary to GAF and as previously shown[2], Zelda is not obviously retained on mitotic chromosomes. Immunostaining of *GFP-zld* embryos confocal images with anti-GFP (green) and anti-GAF (red) are shown at different time points: mitosis between nc12 and nc13 (top row panels), interphase nc13 (second row panels), mitosis between nc13 and nc14 (third row panels) and interphase nc14 (bottom row panels).

(**b**) Immunostaining using anti-GFP and anti-Ser5-P Pol II in *GFP-zld* embryo from the end of mitosis between nc12 and nc13 to nc14 showing that Zelda comes back very quickly in the nucleus at the end of mitosis compared to Ser5-P Pol II. Scale bars represent 10µm.

(**c**) Living *eGFP-bcd/+;His2Av-mRFP/+* embryo imaged by confocal microscopy from interphase of nc13 to early interphase of nc14. Successive representative maximum intensity projected Z-stack images are shown at the indicated timings (see also Supplementary movie 8). Scale bars represent 20µm.

(**d**) Average intensity profiles for nucleoplasmic GFP-Zelda (dark green) and nucleoplasmic eGFP-Bicoid (light green) measured from a nc13 embryo transitioning into nc14. An automatic tracking of fluorescence from a minimum of 87 nuclei generated these profiles, error bars represent SD. Synchronization of the developmental timing was done using the time frame where mother nuclei are splitting into two daughter nuclei, using His2Av-mRFP staining.

(**e**) Example of a time trace obtained by FCS from a *GFP-ash1* nc14 embryo that showed no bleaching. (**e'**) Example of autocorrelation function (black dots) related to (**e**) (red curves represent fitting using diffusion reaction model). (**f**) Box plot representing values for G(∞) (see eq(10)). Centered lines represent the median.

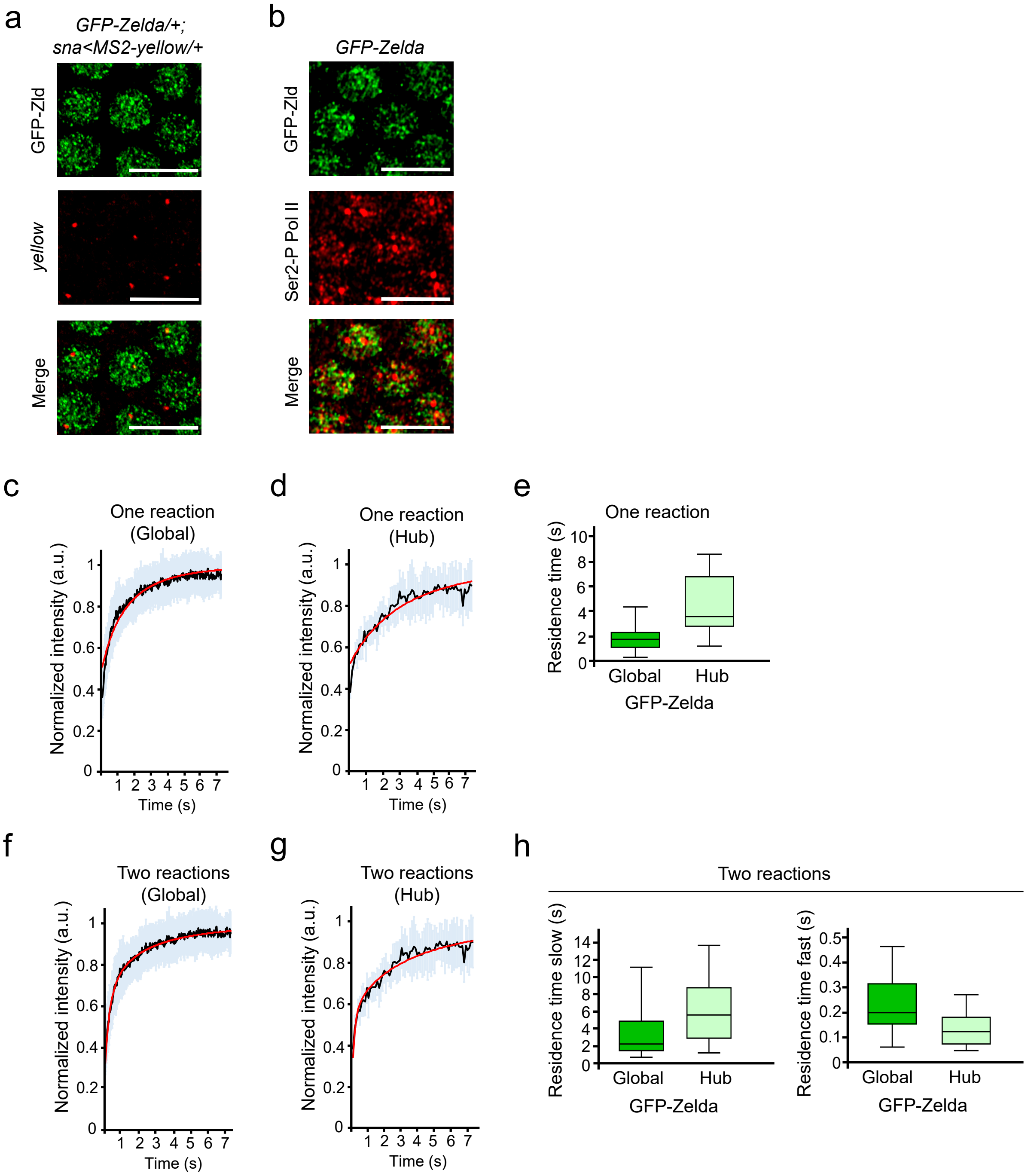


i

| Fitting models | | Residence time Global (s) | Residence time Hub (s) |
|---|---|---|---|
| Reaction only | | 1.9 +/- 1 | 4.3 +/- 2.2 |
| Diffusion reactions | | 3.3 +/- 2.8 | 2.5 +/- 1.6 |
| Two reactions | slow | 3.2 +/- 2.3 | 5.7 +/- 3.7 |
| | fast | 0.23 +/- 0.1 | 0.14 +/- 0.07 |

**Supplementary Figure 6: (related to Figure 6)**
(a) Immuno-FISH using anti-GFP antibody (green) and probe against *yellow* (red) in *GFP-Zld* embryos expressing the *SnaE+1Zld3'* transgenes showing limited colocalization between Zelda hubs and transcription site. Scale bars represent 10µm. (**b**) Immunostaining using anti-GFP (green) and anti-Ser2-P Pol II (red) antibodies in *GFP-Zld* embryos showing no obvious colocalization between Zelda hubs and Pol II. Scale bars represent 10µm. (**c**) Fluorescence Recovery After Photobleaching (FRAP) mean curve (black) and the mean of all the fits (red curve) using one reaction models determined at the bleached spot (Global) for 25 nuclei from nc14 developing *GFP-zld* embryos. The error bars represent SD from different nuclei. (**d**) FRAP mean curve (black) and the mean of all the fit (red curve) using one reaction models determined at the bleached spot (Hub) for 11 nuclei from nc10-13 developing *GFP-zld* embryos. The error bars represent SD from different nuclei. (**e**) Box plot representing the extracted residence time between Global FRAP and Hub FRAP using one reaction models. Centered lines represent the median. (**f**) FRAP mean curve (black) and the mean of all the fits (red curve) using two reaction models determined at the bleached spot (Global) for 25 nuclei from nc14 developing *GFP-zld* embryos. The error bars represent SD from different nuclei. (**g**) FRAP mean curve (black) and the mean of all the fit (red curve) using two reaction models determined at the bleached spot (Hub) for 11 nuclei from nc10-13 developing *GFP-zld* embryos. The error bars represent SD from different nuclei. (**h**) Box plot representing the two extracted residence time (slow and fast) between Global FRAP and Hub FRAP using two reaction models. Centered lines represent the median. (**i**) Table showing the mean residence time +/- SD using the three fitting models for Global or Hub FRAP.

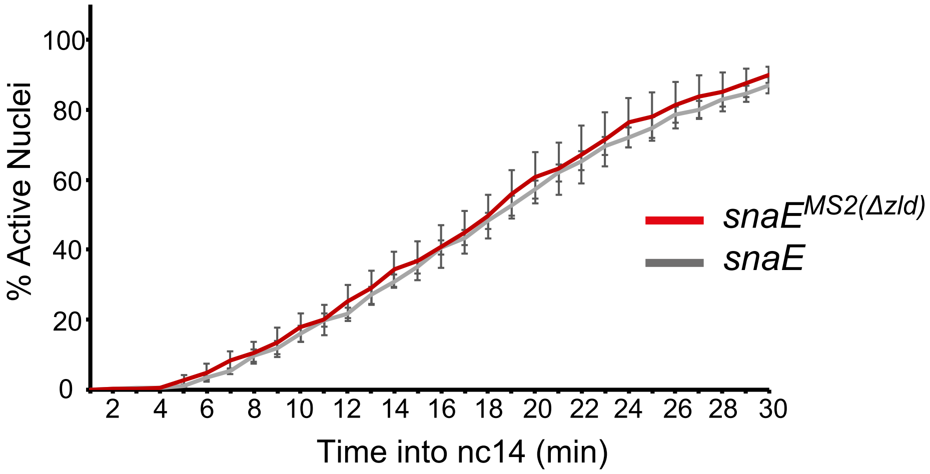


**Supplementary Figure 7:**
Synchrony curve for *snaE* with *MS2deltaZld* (red line)[3] transgenes compared to that of the *snaE* (grey line) transgenes. Statistics: *snaE* (6 movies, n=970 nuclei), *snaE* with *MS2deltaZld* (3 movies, n=456 nuclei). Error bars represent SEM.

**Supplementary Movies:**

Each supplemental movie represents a visual illustration of the data, contrast and brightness were adjusted for better visualization, all movies are a maximum intensity projection of Z-stacks, except for Supplementary Movie 11 and 12.

**Movie 1:** Live imaging of transcriptional activity driven by *snaE*. Nuclei are detected using His2Av-mRFP and MS2 using MCP-GFP.

**Movie 2:** Live imaging of transcriptional activity driven by *snaE+1Zld 5'*. Nuclei are detected using His2Av-mRFP and MS2 using MCP-GFP.

**Movie 3:** Live imaging of transcriptional activity driven by *snaE+2Zld*. Nuclei are detected using His2Av-mRFP and MS2 using MCP-GFP.

**Movie 4:** Live imaging of transcriptional activity driven by *snaE+3Zld*. Nuclei are detected using His2Av-mRFP and MS2 using MCP-GFP.

**Movie 5:** Live imaging of transcriptional activity driven by *snaE+3Zld* in a *white-RNAi* background. Nuclei are detected using His2Av-mRFP and MS2 using MCP-GFP.

**Movie 6:** Live imaging of transcriptional activity driven by *snaE+3Zld* in a *zelda-RNAi* background. Nuclei are detected using His2Av-mRFP and MS2 using MCP-GFP.

**Movie 7:** Living *GFP-zld/+;His2Av-mRFP/+* embryo imaged by confocal microscopy.

**Movie 8:** Living e*GFP-bcd/+;His2Av-mRFP/+* embryo imaged by confocal microscopy.

**Movie 9:** Live imaging of *GFP-zelda* embryo from nc11 to nc14 showing local inhomogeneities.

**Movie 10:** Example of GFP-Zld hubs and their fast movement.

**Movie 11:** Live imaging of Zelda (green) in *GFP-zelda* embryo containing the *snaE+1Zld3'* transgene and the MCP-RFPt (MS2 in Red).

**Movie 12:** Example of Fluorescence Recovery After Photobleaching (FRAP) of one GFP-Zld hub, showing fast movement and fast recovery.

**Supplementary methods:**

**Homogeneous jump model**

In this model, all jumps have the same mean duration, denoted $\tau$. The distribution of the random time $T_r$ depends on $\tau$ and on the number of transitions performed in order to reach the *ON* state. If only one transition is utilized to reach the *ON* state, then $T_r$ is exponentially distributed with the parameter $\tau^{-1}$. Likewise, in the case where the *ON* state is reached after performing $k$ transitions, then $T_r$ is gamma distributed with shape parameter $k$ and with scale parameter $\tau$. In general, $T_r$ is given by a mixture of gamma distributions with shape parameters 1,2,3, ... and scale parameter $\tau$, whose cumulative distribution function (cdf) reads:

$$F_r(t) = P[T_r \leqslant t] = p_1\left(1 - \exp\left(-t/\tau\right)\right) + p_2 \frac{1}{\Gamma(2)}\gamma\left(2, t/\tau\right) + p_3 \frac{1}{\Gamma(3)}\gamma\left(3, t/\tau\right) + \cdots, \quad \text{(S1)}$$

where $p_1$, $p_2$, $p_3$ are the probabilities of one, two, and three jumps, $\Gamma$, $\gamma$ are the complete and incomplete gamma functions, respectively. The mean and the variance of $T_r$ are as such:

$$E[T_r] = (p_1 + 2p_2 + 3p_3 + \ldots)\tau \quad \text{(S2)}$$
$$Var[T_r] = (p_1 + 2p_2 + 3p_3 + \ldots)\tau^2$$

We define the following parameters of the mixed distribution that can be computed empirically from the mean and the variance, the first two moments of the distribution, are as such:

$$b = Var\,[T_r]/E\,[T_r], \qquad a = (E[T_r])^2/Var\,[T_r] \quad \text{(S3)}$$

Using Eq.S2 we find that:

$$b = \tau, \qquad a = p_1 + 2p_2 + 3p_3 + \cdots \quad \text{(S4)}$$

showing that the parameter *'a'* represents the average number of transitions and, equivalently, the mean shape parameter of the mixed gamma distribution, whereas *'b'* is the mean transition time.

In the case of three jumps, the model has three independent parameters $b$, $p_1$, $p_2$. The parameter *'a'* can be computed with (S4) and $p_3$ follows from $p_1+p_2+p_3=1$.

The synchrony of re-activation can be quantified by the time needed for activating half of the population. We denote this time $T_{50}$ and we can compute it as the solution of the equation $F_r(T_{50})=0.5$. Eq.S1 shows that $F_r$ depends on the rescaled argument $t/b$, also on $p_1$, $p_2$. Therefore $T_{50}/b=\psi$ where $\psi$ does not depend on *'b'* but depends on $p_1$, $p_2$. By sampling uniformly the possible values of $p_1$, $p_2$ we showed (see Supplementary Fig. 4) that $\psi$ depends on $p_1$, $p_2$ essentially via *'a'*, and that the following approximate relation holds:

$$T_{50} = b\psi(a), \quad \text{(S5)}$$

where, for the three jumps model, $\psi(a) = 0.085a^4 - 0.78a^3 + 2.65a^2 - 2.81a + 1.56$, and $1 \leq a \leq 3$.

Eq.S5 implies that population activation can be delayed either by increasing *‘b’* (the transition time) or by increasing *‘a’* (the number of transitions).

**Heterogeneous jump model**

In this model, the mean jump durations are heterogeneous. Thus, for a three states model, in order to reach the *ON* state, one can perform a single jump ($A_3 \rightarrow ON$ in mean time $b_3$), two jumps ($A_2 \rightarrow A_3$, then $A_3 \rightarrow ON$, in mean times $b_2$ and $b_3$), or three jumps ($A_1 \rightarrow A_2$, then $A_2 \rightarrow A_3$, then $A_3 \rightarrow ON$, in mean times $b_1$, $b_2$ and $b_3$). For uneven mean jump durations, $b_1 \neq b_2 \neq b_3$, the time to perform two or three jumps is no longer gamma distributed. However, even in this case, it is possible to compute the cdf of $T_r$. To this aim, we consider the continuous time Markov chain $M(t)$, that has four states $A_1$, $A_2$, $A_3$, $A_4$ = *ON*, and three transitions $A_1 \overset{k_1}{\rightarrow} A_2, A_2 \overset{k_2}{\rightarrow} A_3,\ A_3 \overset{k_3}{\rightarrow} ON,$ where the transitions rates are reciprocals of the mean transition times, namely $k_i = b_i^{-1}$, $i$=1,2,3. The *ON* state is absorbing, i.e. once it reaches this state, the process remains there. Then, the activation time $T_r$ results as a solution to the first passage time problem. More precisely, let $T_j = min(t > 0|M(t) = ON, M(0) = A_j)$ be the first passage time in *ON* when $M(t)$ starts from $A_j$, where $j$=1, 2, or 3. If $p_1$, $p_2$, $p_3$ are the probabilities of starting in $A_1$, $A_2$, or $A_3$, respectively, then one has

$$P[T_r \ \leq t] = p_1 P[T_1 \ \leq t] + p_2 P[T_2 \ \leq t] + p_3 P[T_3 \ \leq t] \qquad \text{(S6)}$$

Let $X_{ij}(t) = P[M(t) = A_i|M(0) = A_j]$. Because the state *ON* is absorbing,

$$X_{4j}(t) = \ P\big[M(t) = ON|M(0) = A_j\big] = P[T_j \leq t] \qquad \text{(S7)}$$

Furthermore, because $M(t)$ is a Markov chain, each of the vectors $X_{ij}(t) = (X_{1j}(t), X_{2j}(t), X_{3j}(t), X_{4j}(t))$ satisfies the following system of ordinary differential equations (master equation):

$$\frac{dX}{dt} = QX, \qquad \text{(S8)}$$

where

$$Q = \begin{pmatrix} -\frac{1}{b_1} & 0 & 0 & 0 \\ \frac{1}{b_1} & -\frac{1}{b_2} & 0 & 0 \\ 0 & \frac{1}{b_2} & -\frac{1}{b_3} & 0 \\ 0 & 0 & \frac{1}{b_3} & 0 \end{pmatrix}$$

The solution of Eq.S8 is $X(t) = exp(Qt)X(0)$, where $exp(Qt) = 1 + Qt + (Q^2t^2)/2! + \ldots$ The corresponding initial conditions are $X_1(0) = (1, 0, 0, 0)$, $X_2(0) = (0, 1, 0, 0)$, $X_3(0) = (0, 0, 1, 0)$ and $X_4(0) = (0, 0, 0, 1)$. Next, it can be easily shown that

$$\exp(Qt) = \begin{pmatrix} e^{-\frac{t}{b_1}} & 0 & 0 & 0 \\ \frac{b_2}{b_1-b_2}\left(e^{-\frac{t}{b_1}} - e^{-\frac{t}{b_2}}\right) & e^{-\frac{t}{b_2}} & 0 & 0 \\ \frac{b_1b_2b_3\left(e^{-\frac{t}{b_1}}-e^{-\frac{t}{b_2}}\right)-b_1b_3^2\left(e^{-\frac{t}{b_1}}-e^{-\frac{t}{b_3}}\right)+b_2b_3^2\left(e^{-\frac{t}{b_2}}-e^{-\frac{t}{b_3}}\right)}{(b_1-b_2)(b_1-b_3)(b_2-b_3)} & \frac{b_3\left(e^{-\frac{t}{b_1}}-e^{-\frac{t}{b_2}}\right)}{b_2-b_3} & e^{-\frac{t}{b_3}} & 0 \\ 1-\frac{b_1^2b_2e^{-\frac{t}{b_1}}-b_1b_2^2e^{-\frac{t}{b_2}}+b_1b_3^2e^{-\frac{t}{b_3}}+b_2^3b_3e^{-\frac{t}{b_2}}-b_2b_3^2e^{-\frac{t}{b_3}}}{(b_1-b_2)(b_1-b_3)(b_2-b_3)} & 1-\frac{b_2e^{-\frac{t}{b_2}}-b_3e^{-\frac{t}{b_3}}}{b_2-b_3} & 1-e^{-\frac{t}{b_3}} & 1 \end{pmatrix}. \quad \text{(S9)}$$

Using Eq.S7 and Eq.S9 it follows

$$P[T_3 \leq t] = X_{43}(t) = (0,0,0,1)\, exp(Qt)\, (0,0,1,0)^T = 1 - e^{-\frac{t}{b_3}},$$

$$P[T_2 \leq t] = X_{42}(t) = (0,0,0,1)\, exp(Qt)\, (0,1,0,0)^T = 1 - \frac{b_2e^{-\frac{t}{b_2}} - b_3e^{-\frac{t}{b_3}}}{b_2 - b_3},$$

$$P[T_1 \leq t] = X_{41}(t) = (0,0,0,1)\, exp(Qt)\, (1,0,0,0)^T$$

$$= 1 - \frac{b_1^2b_2e^{-\frac{t}{b_1}} - b_1b_2^2e^{-\frac{t}{b_2}} + b_1b_3^2e^{-\frac{t}{b_3}} + b_2^3b_3e^{-\frac{t}{b_2}} - b_2b_3^2e^{-\frac{t}{b_3}}}{(b_1-b_2)(b_1-b_3)(b_2-b_3)},$$

where $(\ )^T$ stands for transpose.

Finally, using Eq.S6 we obtain

$$F_r(t) = p_1\left(1 - \frac{b_1^2b_2e^{-\frac{t}{b_1}}-b_1b_2^2e^{-\frac{t}{b_2}}+b_1b_3^2e^{-\frac{t}{b_3}}+b_2^3b_3e^{-\frac{t}{b_2}}-b_2b_3^2e^{-\frac{t}{b_3}}}{(b_1-b_2)(b_1-b_3)(b_2-b_3)}\right) + p_2\left(1 - \frac{b_2e^{-\frac{t}{b_2}}-b_3e^{-\frac{t}{b_3}}}{b_2-b_3}\right) + p_3\left(1 - e^{-\frac{t}{b_3}}\right) \quad \text{(S10)}$$

**Modelling memory and memory erasure**

Our model explains the part of memory resulting from differences of *'a'* (number of jumps needed to reach the *ON* state) between descendants of active and inactive mothers. We simulated activation in nc14 by considering that, before mitosis, descendants of active mothers start in $A_3$, whereas descendants of inactive mothers start in $A_1$. During mitosis (5 minutes), the *ON* state is not accessible and downward transitions $A_3 \rightarrow A_2$ and $A_2 \rightarrow A_1$ are possible. After mitosis, the irreversible transition scheme is restored. We have simulated this model and computed the ratio $a_{inactive}/a_{\text{active}}$ corresponding to the number of steps needed to reach the *ON* state after mitosis for nuclei coming from inactive and active mothers, respectively. We noticed that the post-mitotic ratio is smaller than the pre-mitotic value (which is 3), the reduction resulting from mitotic transitions between non-productive states. Furthermore, we computed the dependence of the post-mitotic ratio on the transition time *'b'* and showed that this ratio increases with *'b'*, it is equal to one for small *'b'*, and it is larger than one for large *'b'*. This result is robust with respect to the simulation parameters. The free parameter in the simulation is the ratio of transition times for upward and backward transitions during mitosis. The theoretical curve in the Fig. 4e was obtained when backward transitions are two times slower than upward transitions.

**Data analysis and parameter estimates for modeling**

Data analysis and parameter estimates were performed using MATLAB and Optimization Toolbox Release 2013b, The MathWorks, Inc., Natick, Massachusetts, United States.

The time origin was first set at the end of mitosis. The deterministic waiting time $T_0$ was estimated as the time between the end of mitosis and the time when the first nucleus from a large population is activated. This estimate is accurate for a large number of nuclei, because the probability $p_0$ that $T_r$ is close to zero was supposed to be non-zero. The estimate is more accurate when the number of nuclei is large (by the law of large numbers) and when the number of transitions is small (because $p_0$ is high for a gamma distribution with small shape parameter, as it is the case when the number of transitions is small). Then the origin of time was set at $T_0$ and $T_r$ was determined for all nuclei. Parameters *'a'* and *'b'* were first, roughly estimated with the formulas (S3). We found that *'a'* is not higher than 3, which allowed us to restrict the analysis to only three unidirectional transitions.

The empirical cumulative distribution function of $T_r$ was estimated using the Kaplan-Meier method. The homogeneous jump model Eq.S1 and the heterogeneous jump model Eq.S10 were fitted by minimization of an objective function *O* defined as the $l^2$ (sum of squares) distance between experimental data and model prediction. The optimization was performed using the MATLAB function *lsqnonlin* starting from 100, different, randomly chosen initial parameter values. The best optimum and the suboptimal solutions where the objective function was at most 50% higher than the best optimum were used to quantify parameter uncertainty. For instance, if the optimal and suboptimal values of the parameter *p* are between $p_{min}$ and $p_{max}$, its uncertainty is computed as

$$err_p = \frac{p_{max}-p_{min}}{p_{max}+p_{min}} \quad \text{(S11)}$$

The results of the fit are the three parameters $p_1$, $p_2$, $p_3$ and *'b'* (Supplementary Tables 1). The parameter *'a'* is also computed using Eq.S4. The values of *'a'* and *'b'* estimated from Eq.S3 and from the distribution fit can be different. The first method, based on the moments, is sensitive to extreme values; therefore, it is less reliable than the latter.

The results of the parameter estimation are given in the Supplementary Tables 1 and 2.

For Fig. 4c-e and Supplementary Tables 1 and 2, after checking that single genotype estimates were equivalent to the pooled data, we decided to present the estimates for *'a'* population of nuclei pooled from *Zelda-RNAi snaE* and *Zelda-RNAi snaE+1Zld 5'* embryos. Given that the only difference between *snaE* and *snaE+1Zld 5'* transgenes resides in the added CAGGTAG, we reasoned that in the absence of Zelda, these genotypes are theoretically comparable, thus justifying their pooling, to increase sample size.
